# Identification of a phage display-derived peptide interacting with the N-terminal region of Factor VII activating protease (FSAP) enables characterization of zymogen activation

**DOI:** 10.1101/2022.01.09.475526

**Authors:** Sebastian Seidl, Nis Valentin Nielsen, Michael Escheid, Bengt Erik Haug, Maria Stensland, Bernd Thiede, Paul J. Declerck, Geir Åge Løset, Sandip M. Kanse

**Affiliations:** Oslo University Hospital and Medical Faculty, University of Oslo, Oslo, Norway; Paul Ehrlich Institute, Langen, Germany; Department of Chemistry and Center for Pharmacy, University of Bergen, Bergen, Norway; Department of Biosciences, University of Oslo, Oslo, Norway; Department of Pharmaceutical and Pharmacological Sciences, Katholieke Universiteit Leuven, Belgium; Nextera, Oslo, Norway

**Author notes:** Joint 1^st^ authors (Sebastian Seidl, Nis Valentin Nielsen). **Correspondence:** Sandip M. Kanse, Institute for Basic Medical Sciences, University of Oslo, 0172, Oslo, Norway.

**Keywords:** FSAP, HABP2, zymogen activation, peptide, phage display, hemostasis

## Abstract

Increased Factor VII activating protease (FSAP) activity has a protective effect in diverse disease conditions as inferred from studies in FSAP^−/−^ mice and humans deficient in FSAP activity due to a single nucleotide polymorphism. The activation of FSAP zymogen in plasma is mediated by extracellular histones that are released during tissue injury or inflammation or by positively charged surfaces. However, it is not clear if this activation mechanism is specific and amenable to manipulation. Using a phage display approach we have identified a peptide, NNKC9/41, that activates pro-FSAP in plasma. Other commonly found zymogens in the plasma were not activated. Binding studies with FSAP domain deletion mutants indicate that the N-terminus of FSAP is the key interaction site of this peptide. Blocking the contact pathway of coagulation did not influence pro-FSAP activation by the peptide. In a monoclonal antibody screen, we identified MA-FSAP-38C7 that prevented the activation of pro-FSAP by the peptide. This antibody bound to the LESLDP sequence (amino acids 30-35) in the N-terminus of FSAP. The plasma clotting time was shortened by NNKC9/41 and this was reversed by MA-FSAP-38C7 demonstrating the utility of this peptide. Identification of this peptide, and the corresponding interaction site, provides proof of principle that it is possible to activate a single protease zymogen in blood in a specific manner. Peptide NNKC/41 will be useful as a tool to delineate the molecular mechanism of activation of pro-FSAP in more detail, elucidate its biological role.

**Graphical abstract:** 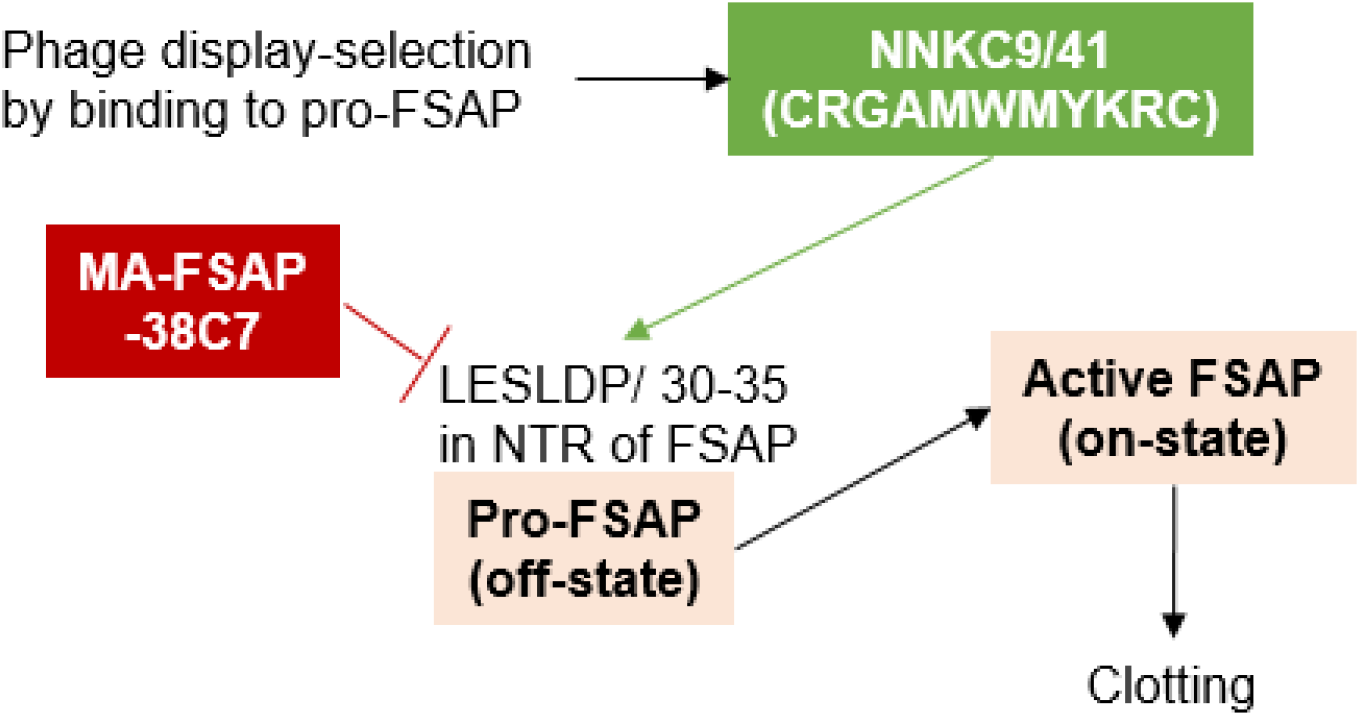

## INTRODUCTION

Factor VII activating protease (FSAP) is a serine protease, predominantly, expressed in the liver and encoded by the hyaluronic acid binding protein2 (*HABP2*) gene. A naturally occurring mutation (Gly534Glu) renders the protease inactive ^*1*^ and this single nucleotide polymorphism (SNP), termed Marburg I (MI) SNP, is present in 5% of the Caucasian population. It is associated with enhanced risk of stroke ^*2*^, carotid stenosis ^*3*^ and venous thrombosis ^*4, 5*^. These conclusions are based on a candidate-gene approach and are contentious in the case of venous thrombosis ^*6, 7*^. FSAP^−/−^ mice show increased liver fibrosis ^*8*^, stroke ^*9*^ and neointima formation ^*10*^ compared to their wild type counterparts, but the extent of thrombosis is lower ^*11*^. Hence, a lower FSAP activity is linked to an increased disease burden in both mice and humans. This led us to hypothesize that increasing the FSAP activity in plasma will have a positive therapeutic effect in some conditions e.g., stroke. This could be achieved, for instance, by administering recombinant FSAP or by activating endogenous pro-FSAP.

Pro-FSAP circulates as a single chain inactive zymogen in the plasma. Negatively charged polymers such as heparin ^*12, 13*^, nucleic acids ^*14, 15*^ and dextran sulfate ^*12*^ activate purified pro-FSAP to various degrees. Positively charged histones ^*16*^ as well as positively-charged surfaces ^*17*^ activate pro-FSAP in blood, plasma and *in vivo* ^*16, 17*^. The activation of pro-FSAP is induced by the interaction of anions or cations with the N-terminal region (NTR) or the EGF3 domain of FSAP, respectively. Yamamichi et al have suggested that charged molecules induce a conformational change, promoting dimerization of pro-FSAP molecules. This leads to cleavage of the interacting partner molecule at the activation site Arg313-Ile314 leading to full activity ^*18*^.

*In vivo*, FSAP activity is increased after surgery and in patients with sepsis ^*19*^, trauma ^*20*^ and acute respiratory distress syndrome ^*21*^ which suggests that pro-FSAP activation is related to tissue damage and inflammation. Histones have ubiquitous effects in plasma, blood and cells ^*22, 23*^ and the activation of FSAP may be part of their function as damage associated molecular pattern (DAMPs). Substrates of active FSAP include proteins from the hemostasis and complement system as well as growth factors, histones ^*24*^ and protease-activated receptors (PARs) ^*25*^.

Currently, it is not known if the activation of pro-FSAP is simply dependent on charged macromolecules or if there is a higher specificity to this process. Phage display has been successfully used as a strategy to identify peptides that activate the zymogen form of proteases ^*26*^. We hypothesized that a phage display screen could be used to identify peptidic modulators of pro-FSAP activity. This approach led to the identification of a peptide that can activate pro-FSAP in a specific, selective and potent manner. Furthermore, we identified an inhibitory antibody that blocks pro-FSAP activation by the peptide and demonstrate the efficacy of the peptide and the antibody in plasma clotting assays. Thus, we have established that the activation of pro-FSAP is a precise and exploitable mechanism.

## RESULTS and DISCUSSION

### Selection of pro-FSAP binding phages

To identify peptidic binders and modulators of pro-FSAP activity, we designed a peptide phage display approach to selectively enrich for strong binders using phage selection against biotinylated pro-FSAP purified from human plasma. Two parallel approaches were chosen, where either a random 11-mer (NNK11) or a Cys-constrained random 9-mer (NNKC9) peptide library (formally an 11-mer) were panned in three iterative rounds with decreasing amount of bead-immobilized target protein in each round. The selection was tuned to favor binders with high affinity by altering the peptide display levels from high to low valency in later selection rounds as well as using excess pro-FSAP as a soluble competitor to deplete low affinity binders in all three rounds (Fig. 1A). We then screened phage clones from the third selection round of each library in a phage ELISA-based pro-FSAP binding assay. From the NNKC9 library, 5/6 positive binding clones showed the sequence CRGAMWMYKRC, termed NNKC9/41 and the remaining clone had the sequence CEGLAIQVKQC (Fig. 1A). From the NNK11 library, 10/11 positive binding clones had the sequence IDCLMQNAGSA, termed NNK11/189, and the remaining clone had the sequence DLPWSMPRPCR (Fig. 1A). Thus, we identified two predominant sequences capable of binding to pro-FSAP (sequences in Fig. 1A and Supplementary Table 1) that were investigated further.

**Fig. 1.**
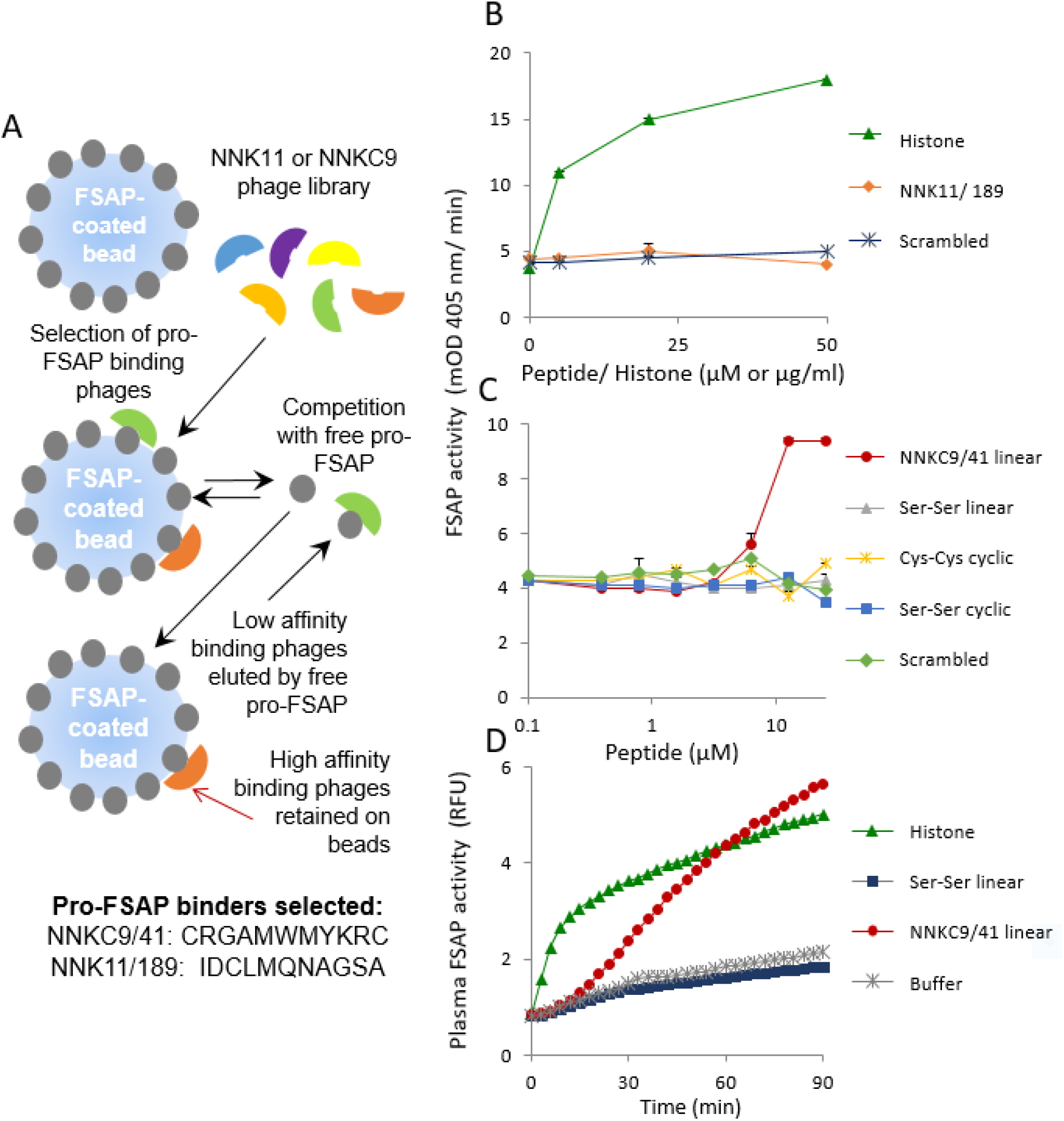
Isolation of pro-FSAP binding peptides and their activity: (A) The strategy to isolate high affinity phages from the NNKC9 and NNK11 library that bind to pro-FSAP. To select for strong binders an excess of free pro-FSAP protein was used for competitive elution of low affinity binders. High affinity binders retained on the beads were analyzed further. The sequences of major positive clones identified through an ELISA-based binding assay from the NNKC9 and the NNK11 library are shown. (B) NNK11/189 and scrambled controls (0-50 μM) as well as histones (0-50 μg/ml) were added to purified pro-FSAP (1.0 μg/ml) and the turnover of the chromogenic substrate S2288 (200 μM) was measured. (C) Linear NNKC9/41 was compared to a peptide that was synthetically cyclized (Cys-Cys cyclic), peptide with Ser residues instead of Cys at both ends (Ser-Ser linear) and peptide with Ser residues at the end cyclized head to tail (Ser-Ser cyclic) as well as a scrambled peptide in a pro-FSAP activation assay. (D) Hirudin plasma (1:12 dilution) was incubated with histones (25 μg/ml), NNKC9/41 or Ser-Ser linear control (each, 25 μM) and the fluorogenic substrate turnover (Ac-Ala-Lys-Nle-Arg-AMC) was monitored in duplicate wells and results from a single well are shown. In panels B-C, results are shown as mean + range (duplicate wells). Results were replicated with 5 different batches of synthetic linear NNKC9/41 peptide (panels B-C) and on plasma from 5 different donors (panel D). Activity in the absence of peptide or histones (panels B, C) is due to the presence of active FSAP in the preparation of pro-FSAP.

### Activation of pro-FSAP by NNKC9/41 but not NNK11/189

To test the effects of these synthetic peptides we used pro-FSAP, purified from plasma, and measured its activation or inhibition of its activation using a chromogenic substrate (S2288). Histones were used as a positive activator in all these experiments. NNK11/189, or its scrambled control (see Supplementary Table 1 for sequences), did not have any effect on pro-FSAP activation compared to a strong effect of histones (Fig. 1B). NNKC9/41, synthesized with an internal disulfide bond (NNKC9/41 cyclic) showed no effect. However, the linear peptide with reduced Cys was able to robustly activate pro-FSAP (Fig. 1C). We then assessed the importance of the Cys residues by replacing them with Ser residues. A linear peptide with two Ser residues (Ser-Ser linear) as well as the head-to-tail cyclized version (Ser-Ser cyclic) failed to activate pro-FSAP (Fig. 1C). Scrambling the NNKC9/41 peptide sequence also led to a complete loss of activity (Fig. 1C). Thus, the Cys residues, as well as their positioning, were important for the activation of pro-FSAP. These results were replicated with 5 different batches of the synthetic linear peptide and 4 batches of circular peptide from two different sources indicating that the activity was stable and reproducible.

The next question was if the linear NNKC9/41 peptide could activate pro-FSAP in human plasma which is a complex environment. Since the chromogenic substrate is not suitable for plasma experiments we used the fluorescent substrate (Ac-Ala-Lys-Nle-Arg-AMC) ^*24*^. Linear NNKC9/41, but not the control Ser-Ser linear peptide, induced a robust activation of pro-FSAP comparable to that with histones (Fig. 1D). These results were replicated in plasma from 5 donors. The degree of pro-FSAP activation was variable, presumably reflecting the concentration of FSAP antigen and ambient inhibitor concentration in each donor plasma (data not shown).

It is believed that Cys-constrained M13 phage peptide libraries, e.g., NNKC9, primarily display cyclic peptides as the capsids are assembled into the virions in the oxidized periplasma of *E. coli* ^*27*^. Paradoxically, the synthetic Cys-cyclic peptide was inactive in the assay. However, the presentation of the peptide on the phage versus the synthetic peptide in solution may differ. The activity of the linear peptide may be explained by other factors. E.g., a peptide with Cys residues at both ends can not only form a cyclic monomer but also several cyclic or linear multimeric forms through head-to head, tail-to-tail or head-to-tail disulphide bond formation. Some peptide sequences also have a propensity to form non-covalent aggregates ^*28*^.

To resolve this issue we performed RP-HPLC and liquid chromatography-mass spectrometry (LC-MS) analysis on linear NNKC9/41. This confirmed the sequence identity of the synthesized peptide and the activity of the re-purified peptide. Interestingly, when the cyclic peptide was reduced with DTT, repurified by RP-HPLC and tested, robust activation of pro-FSAP in plasma was observed (data not shown) (Supplementary Fig, 1 and 2). LC-MS analysis revealed the presence of cyclic, di-cyclic peptides with varying degree of oxidation in the linear synthetic NNKC9/41 preparation (Supplementary Fig. 1 and 2). We are currently using diverse purification strategies to isolate the active moiety to homogeneity and define it more precisely.

### Binding of NNK11/189, NNKC9/41 to pro-FSAP

Next, we examined the binding properties of the peptides to pro-FSAP and other plasma proteins. N-terminally biotinylated NK11/189 and linear NNKC9/41 bound to immobilized pro-FSAP in a concentration-dependent manner whereas their scrambled counterparts exhibited low binding (Fig. 2A). NNKC9/41 did not bind to a selection of plasma proteins at lower peptide concentrations, although some binding was noticeable at concentrations >10 μM (Fig. 2B). At these high concentrations there was a decrease in peptide binding to pro-FSAP for unknown reasons (Fig. 2B).

**Fig. 2.**
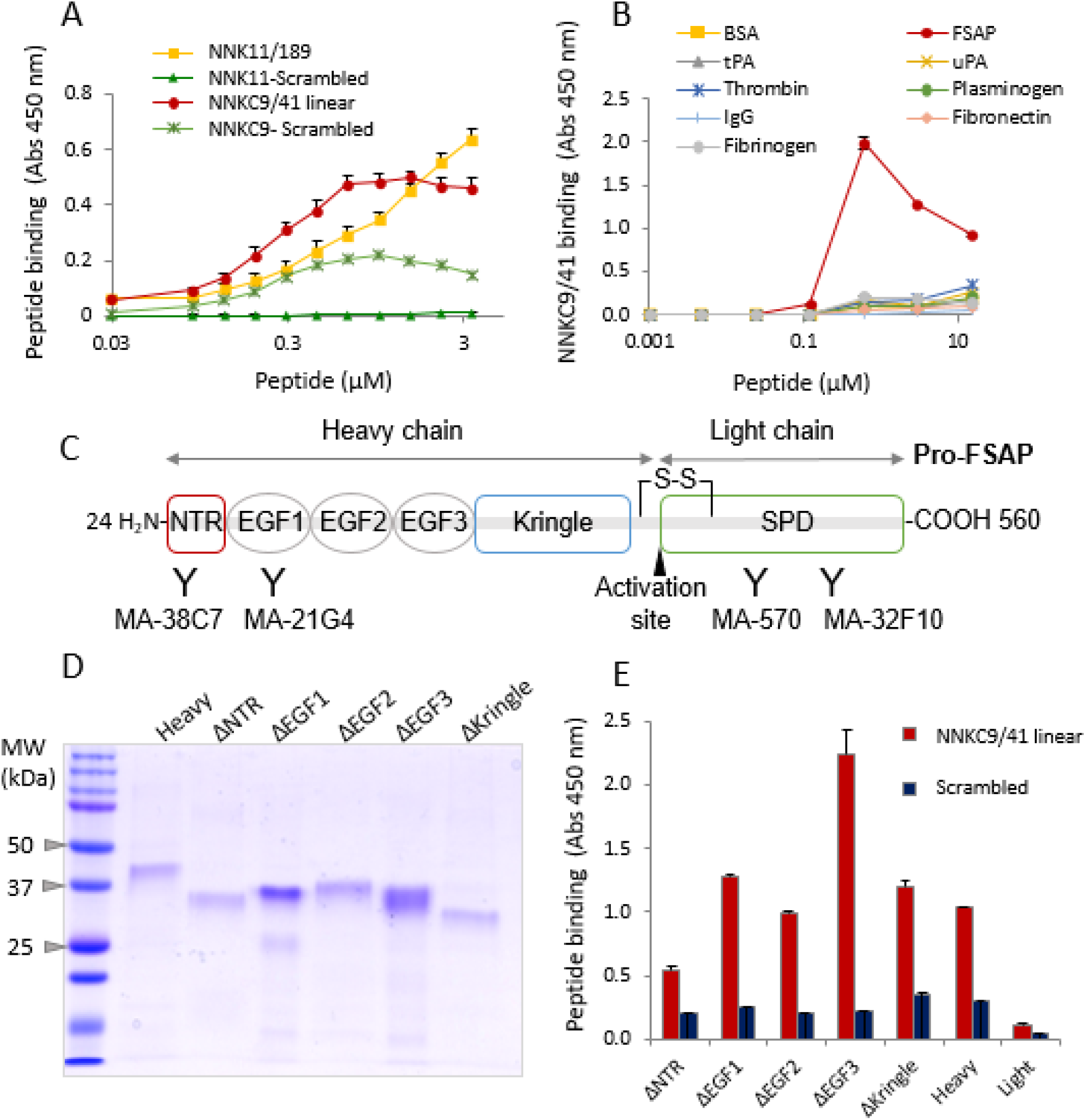
NNKC9/41 interacts with the NTR of pro-FSAP: (A) To determine peptide binding to pro-FSAP, wells were coated with 2μg/ml pro-FSAP and biotinylated NNKC9/41, NNK11/189 and their respective scrambled controls were added in different concentration (0-5 μM). Peroxidase-labelled streptavidin was used to detect peptide binding. (B) Same procedure as in (A) except that different proteins were immobilized and biotinylated NNKC9/41 (0-30 μM) was added and its binding was measured. (C) Schematic structure of pro-FSAP with the heavy chain consisting of an NTR, 3 EGF domains and a kringle domain, followed by the light chain harbouring the serine protease domain (SPD). The binding domains of antibodies used in this study are indicated. (D) Recombinant complete heavy chain (Heavy) and truncated heavy chain variants ΔNTR, ΔEGF1, ΔEGF2, ΔEGF3, and ΔKringle were expressed in bacteria and analyzed by SDS-PAGE followed by Coomassie staining. (E) Wells were coated with 2μg/ml of rabbit polyclonal anti-FSAP antibody and recombinant proteins (1μg/ml) were captured. Biotinylated NNKC9/41 or scrambled control peptide was added at 1 μM and the peptide binding measured with peroxidase-linked streptavidin. In panels A, B and E, results are shown as mean + SEM (triplicate wells).

To identify the domain of FSAP responsible for binding to NNKC9/41, we expressed various domain-deletion mutants of pro-FSAP as His-tagged proteins in *E. coli*. The N-terminal end of pro-FSAP (heavy chain) consists of the unstructured N-terminal region (NTR), three EGF domains and a kringle domain, whereas the C-terminal-light chain contains the serine protease domain (SPD) (Fig. 2C). Histones have been shown to interact with the NTR of FSAP ^*16*^ and heparin binds to the EGF3 domain^*29*^. The recombinant proteins (Fig. 2D) were captured on polyclonal FSAP antibody-coated wells and were used for binding studies. NNKC9/41 exhibited high binding to the heavy chain and very low binding to the light chain/ serine protease domain (SPD) and the ΔNTR mutant (Fig. 2E). The scrambled peptide showed very low binding to all mutants (Fig. 2E). The higher binding of the peptide to the ΔEGF3 mutant could be because this domain, when present, exerts an inhibitory influence on binding. Thus, the binding site for the peptide was in the NTR of pro-FSAP.

We then used another complementary approach to narrow down the site of NNKC9/41 binding to pro-FSAP. For this, we took advantage of a previously generated panel of monoclonal antibodies ^*30*^ and then expanded the screen to identify antibodies that putatively modulated the effect of the peptide on pro-FSAP activation. Indeed, we identified an antibody, MA-FSAP-38C7 which completely blocked the activation of pro-FSAP by NNKC9/41 (Fig. 3A), whereas a control antibody, MA-32F10, had no effect. These experiments were performed with the fluorescent substrate (Ac-Ala-Lys-Nle-Arg-AMC) as opposed to S-2238 in Fig. 2C. MA-FSAP-38C7 did not influence the activity of FSAP if it was already in an activated state (data not shown). Thus, MA-FSAP-38C7, which is directed to the NTR, completely blocked pro-FSAP activation by the peptide (Fig. 2C and 3B).

**Fig. 3:**
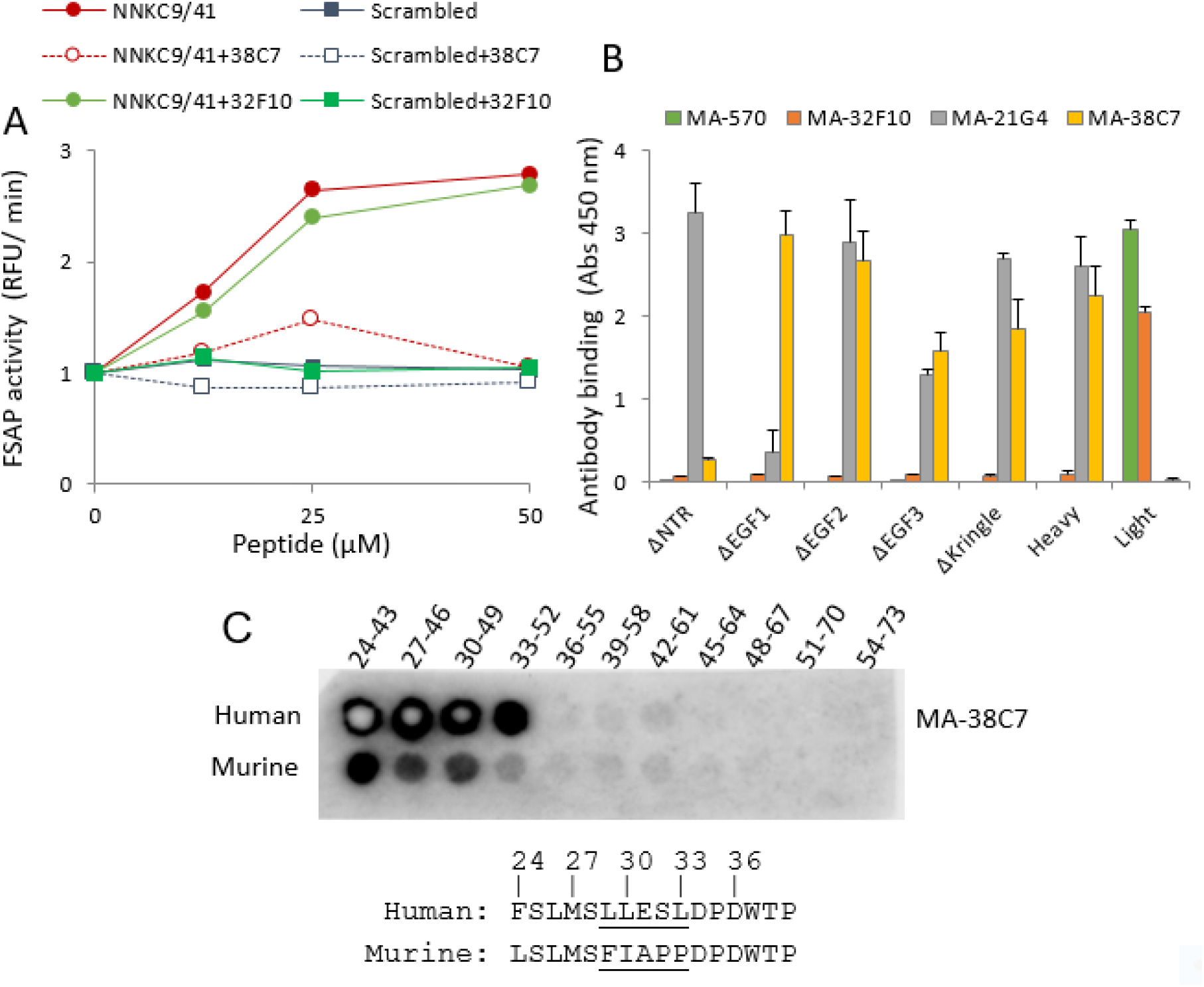
Inhibition of pro-FSAP activation by MA-FSAP-38C7 and identification of its binding epitope: (A) Antibodies (6 μg/ml) were added to pro-FSAP (1.0 μg/ml) followed by NNKC9/41 linear or its scrambled control (0-50 μM each) and activation was determined with fluorogenic substrate Ac-Ala-Lys-Nle-Arg-AMC (80 μM) in duplicate (mean + range). (B) Wells were coated with rabbit polyclonal anti-FSAP antibody (5 μg/ml) and recombinant proteins (2 μg/ml) were captured. The indicated monoclonal antibodies were added at a concentration of 1μg/ml and their binding measured with anti-mouse peroxidase-labelled secondary antibody in triplicates (mean + SEM). (C) 20-mer peptides with a three amino acid shift from the NTR (24-73) of human and mouse pro-FSAP were synthesized on nitrocellulose membranes (University of Oslo, Peptide synthesis Facility). Membranes were blocked with 5% (v/v) BSA and the binding of MA-FSAP-38C7 was detected with appropriate peroxidase-coupled reagents. The MA-FSAP-38C7 binding region in pro-FSAP is amino acids 30-35 and the difference between human and mouse sequence is underlined.

We then used overlapping peptide arrays from the NTR of pro-FSAP to identify the binding site for MA-FSAP-38C7. MA-FSAP-38C7 bound specifically to the LESLDP (amino acids 30-35 in full-length FSAP) sequence in the NTR region. (Fig. 3C). Thus, the LESLDP sequence appears to be important for the activation of pro-FSAP. The murine sequence, which is divergent at this location, exhibited less antibody binding. Biotinylated NNKC9/41 exhibited no specific binding to this peptide array indicating a more elaborate binding interface requiring complex structural elements of the folded protein.

### Pro-FSAP activation by NNKC9/41 in human plasma

Next we characterized the effects of the peptide of pro-FSAP activation in plasma using a multitude of techniques. Our initial studies showed that NNKC9/41 activated pro-FSAP in plasma anti-coagulated with hirudin at plasma dilutions ranging from 1:2 to 1:12 (Fig. 1D and data not shown). In comparison to histone, a much longer lag time in pro-FSAP activation was observed with the peptide (> 15 min) (Fig. 1D). However, in plasma anticoagulated with citrate, the effect of NNKC9/41 was weak at lower dilution of plasma (1:2). Thus, Ca^2+^ is necessary for optimal pro-FSAP activation by the peptide. Western blotting of hirudin, citrate-plasma, after activation with peptides, showed the emergence of a FSAP-inhibitor complex band that is a proxy marker for the activation of pro-FSAP in plasma (Supplementary Fig. S3). This provides additional evidence for peptide-mediated activation of pro-FSAP in plasma.

In order to consolidate the results and exclude any artefacts, we used two additional methods to demonstrate that NNKC9/41 activated pro-FSAP in plasma; (i) Immunocapture of pro-FSAP from plasma, its activation by the peptide and the conversion of the substrate pro-uPA to uPA and (ii) the formation of FSAP-alpha2-antiplasmin complexes in plasma as a readout for pro-FSAP activation and its subsequent inhibition. A concentration dependent effect of NNKC9/41 and histones demonstrated that the peptide was active in all the different tests used to measure pro-FSAP activation in plasma (Fig, 4A-C). The dynamics of the activation process and the properties of the different assays probably account for the fact that the peptide has a stronger effect than histones on fluorogenic substrate turnover Fig. 4A) but is weaker than histones in the other two assays (Fig. 4B-C).

**Fig. 4.**
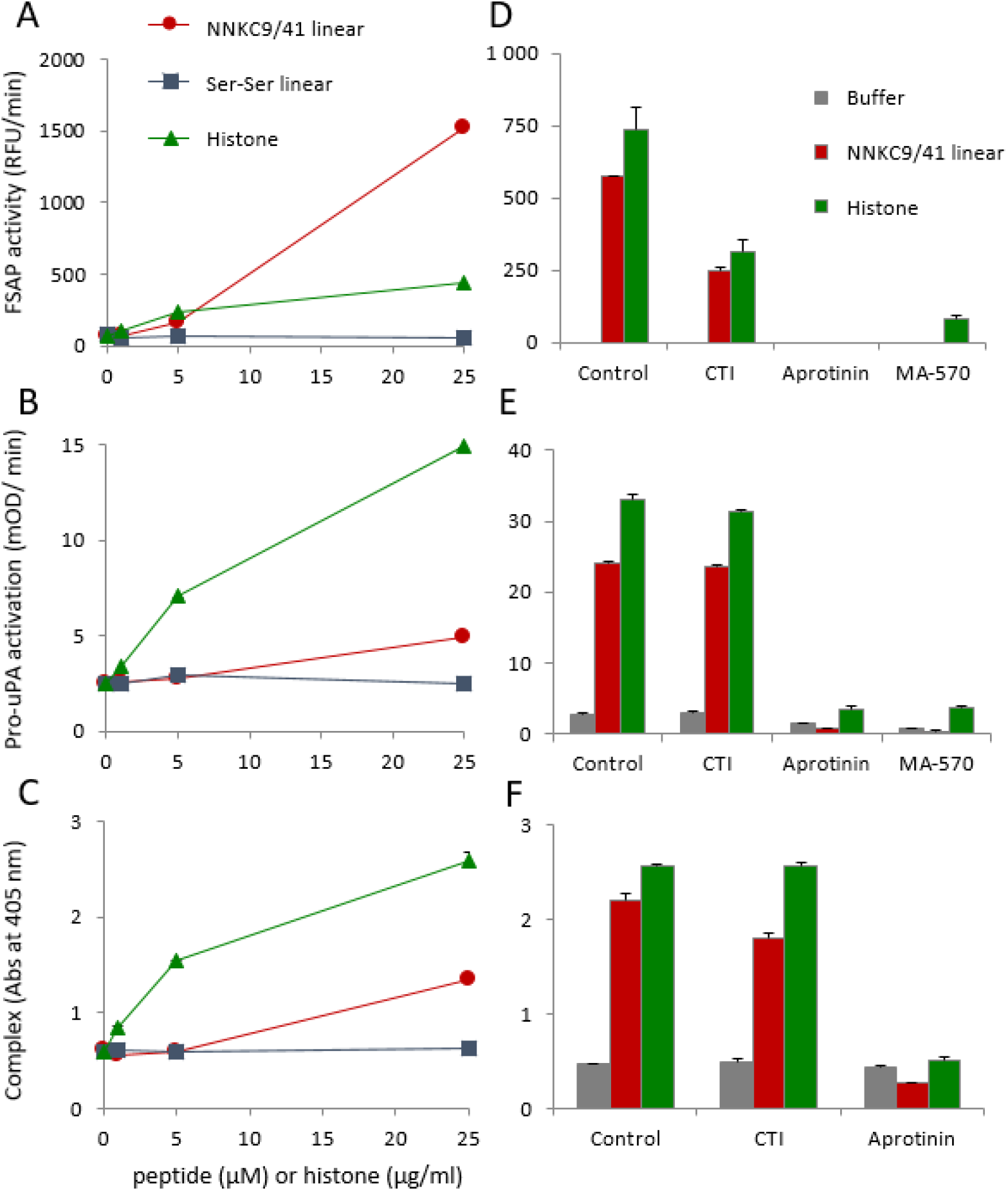
Comparison of different methods to measure pro-FSAP activation in plasma: (A) Hirudin plasma (1:12 dilution) was incubated with histones (0-25 μg/ml), NNKC9/41 linear or its Ser-Ser linear control (0-25 μM) and the fluorogenic substrate turnover (Ac-Ala-Lys-Nle-Arg-AMC) was monitored. (B) From the same experiment as in (A) after 1 h incubation at 37 ºC, samples were captured on anti-FSAP coated wells and the conversion of added pro-uPA to active uPA was measured by determining the turnover of the chromogenic substrate PNAPEP1344 (25 μM). (C) From the same experiment as in (A) after 1 h incubation at 37 ºC, samples were captured on anti-FSAP coated wells and α2-antiplasmin antibody was used to detect the formation of FSAP-α2 antiplasmin complexes. (D, E and F) Hirudin plasma (1:12 dilution) was activated with histones (20 μg/ml) or NNKC9/41 linear (25 μM) in the presence of CTI (80 μg/ml), aprotinin (100 μg/ml) or MA-570 (20 μg/ml) followed by analysis as described in A, B and C respectively. In F, the samples with MA-FSAP-570 were not analyzed since this antibody would interfere in the sandwich ELISA. For panels A-F results are shown as mean + range of duplicates wells and the results were replicated in plasma samples from 3 donors.

We also tested if the contact pathway was involved in the pro-FSAP activation process. Inhibition of the contact pathway by the FXIIa inhibitor, corn trypsin inhibitor (CTI), did not influence the activation of pro-FSAP by the peptide (Fig. 4D-F). The kallikrein inhibitor PKS1-527 had no effect (data not shown) but aprotinin, a general serine protease inhibitor, blocked pro-FSAP activation/ activity (Fig. 4D-F). MA-FSAP-570, a known inhibitory antibody against FSAP also blocked the activity/ activation of FSAP/pro-FSAP (Fig. 4D-E). The inhibitory effect of CTI (80 μg/ml) is most probably due to its direct inhibition of FSAP activity (data not shown). Hence, we can exclude the involvement of the contact pathway in the pro-FSAP activation by the peptide.

The next question was if the peptide was activating a mechanism common to many zymogens ^*26*^ or if it was selective for pro-FSAP. Linear NNKC9/41 did not influence the activity of plasminogen, pro-urokinase, Factor XII, pro-thrombin and Factor X as well as their respective active enzymes (Supplementary Fig. S4). This is also in line with the observation that it did not bind to any other plasma protein tested, except FSAP (Fig. 2B). Thus, the peptide showed very high selectivity for pro-FSAP activation. Linear NNKC9/41 did not alter the activity of the recombinant serine protease domain (SPD) of FSAP (Supplementary Fig. S4A). These changes are consistent with the zymogen activation mechanism rather than an allosteric effect of the peptide on, already, active FSAP.

In the hemostasis cascade, zymogen activation often requires the association of the various components on a macromolecular surface. To mimic this surface-dependent process, we immobilized biotinylated linear NNKC9/41 on neutravidin-coated plates. After capture and preincubation with hirudin plasma, a robust activation of pro-FSAP was observed but this was not the case with the biotinylated scrambled peptide (Fig. 5A). In a corollary of the above experiment, pro-FSAP was captured from plasma on FSAP antibody-coated wells and NNKC9/41 linear and its scrambled counterpart were added to the wells to activate immobilized pro-FSAP. Only NNKC9/41 was found to activate pro-FSAP (Fig 5A), confirming the activity of this peptide in different assay configurations, also with N-terminal biotinylation.

**Fig. 5:**
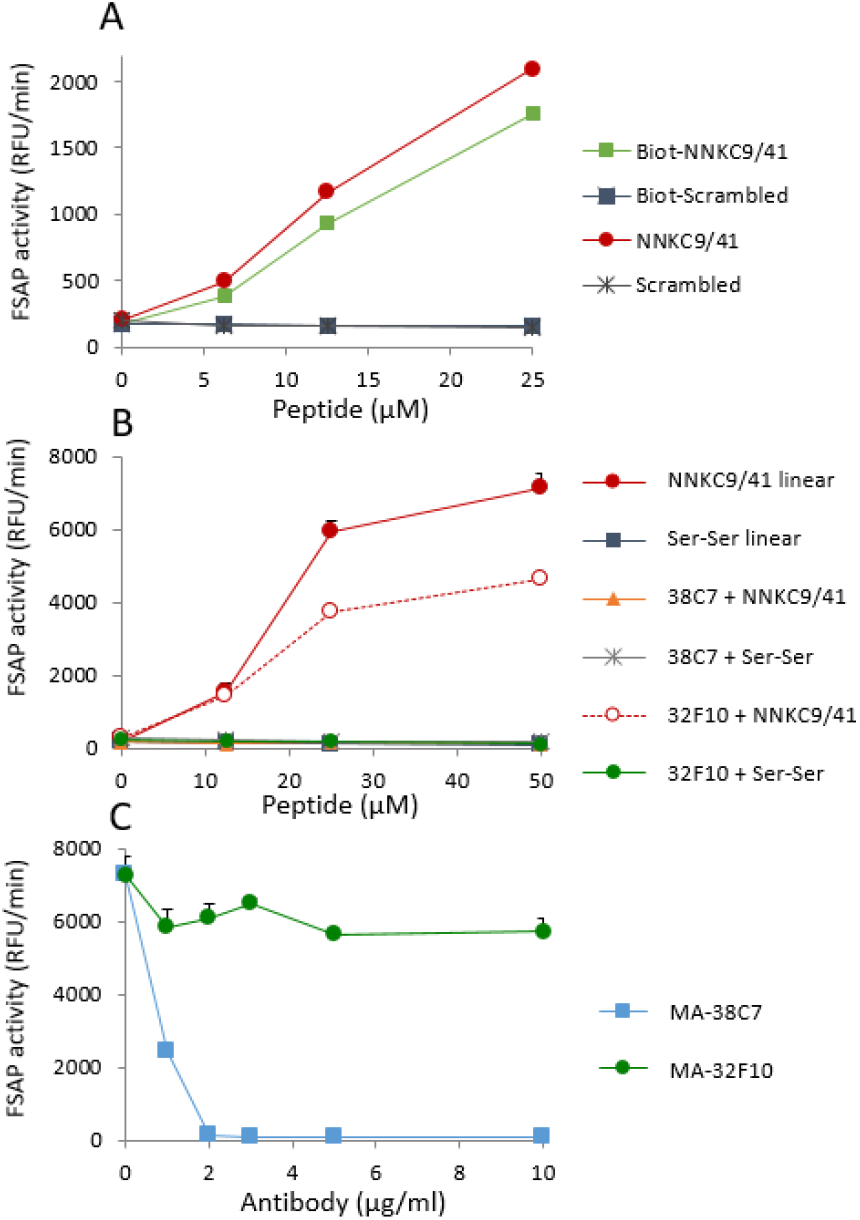
Characteristics of pro-FSAP activation in plasma by NNKC9/41: (A) Neutravidin-coated wells were used to capture biotinylated NNKC9/41 linear or its scrambled control peptide (0-25 μM). Hirudin plasma (1:10) was incubated in the wells for 1 h at 37 ºC to allow for binding and activation of pro-FSAP. In parallel, wells were coated with rabbit polyclonal anti-FSAP antibody (2 μg/ml) and plasma was added to immobilize FSAP in the presence of solution phase NNKC9/41 linear or its scrambled control (0-25 μM). Fluorogenic substrate turnover (Ac-Ala-Lys-Nle-Arg-AMC) was monitored. (B) Plasma was incubated with NNKC9/41 linear and Ser-Ser control (0-50 μM each) in the absence of any antibody or in the presence of MA-FSAP-38C7 or MA-FSAP-32F10 (5 μg/ml) and the FSAP activity was monitored. (C) Plasma was incubated with NNKC9/41 linear peptide (25 μM) in the presence of MA-FSAP-38C7 or MA-FSAP-32F10 (0-10 μg/ml) and the FSAP activity was monitored. For A-C results are shown as mean + range of duplicate wells.

The antibody MA-FSAP-38C7 was then tested for its ability to modulate pro-FSAP activation in plasma. Dose-response analysis showed that MA-FSAP-38C7 antibody was a potent inhibitor of the peptide-mediated activation of pro-FSAP in plasma (Fig. 5B, C), whereas MA-FSAP-32F10 showed a much weaker effect. These antibodies as well as their Fab fragments showed the same effect when histones were used as activators of pro-FSAP (Supplementary Fig. S5).

To consolidate the above results we also performed studies on FSAP-depleted plasma and plasma from persons who were heterozygous or homozygous for the MI-SNP (all anticoagulated with citrate and diluted 1:12). In FSAP-depleted plasma, no turnover of substrate was observed in the presence of the NNKC9/41 peptide. Similarly, the peptide mediated high substrate turnover in WT/WT plasma, low turnover in WT/MI plasma and none in MI/MI plasma (Fig. 6A). These results further underscore the direct, selective and specific effect of the peptide on the activation of endogenous pro-FSAP in plasma.

**Fig. 6:**
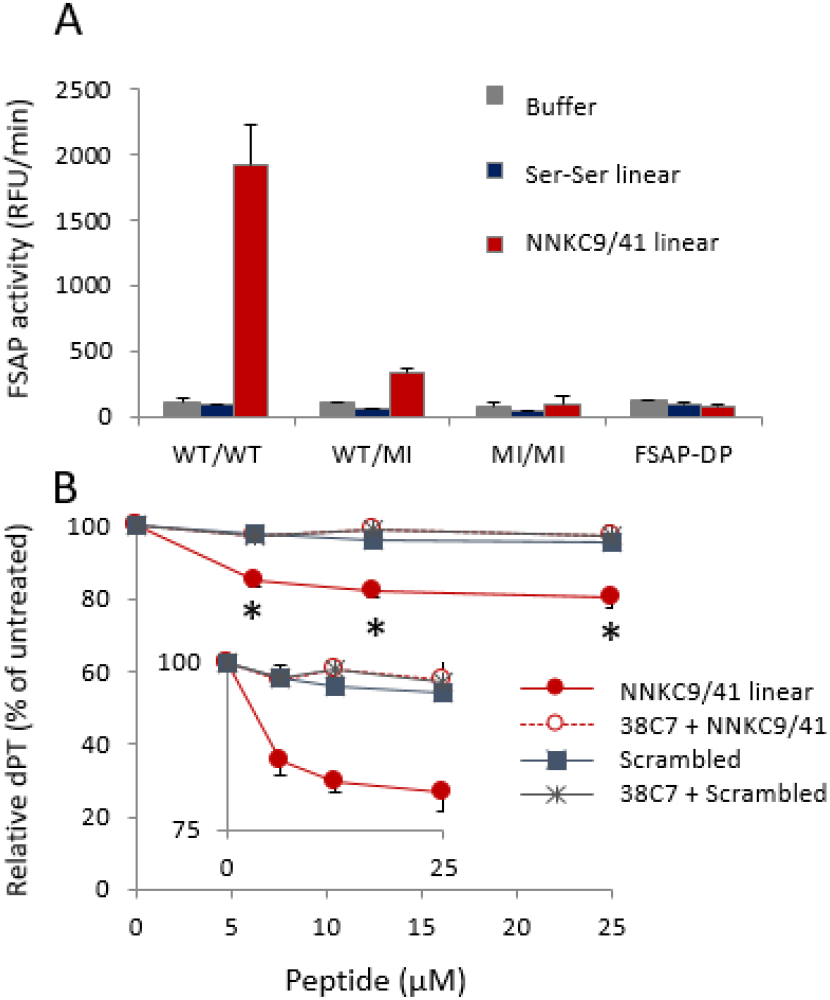
Pro-FSAP activation and clotting in plasma by NNKC9/41: (A) Citrate plasma (1:12 dilution) from donors with WT/WT, WT/MI and MI/MI genotype as well as WT-FSAP-deficient plasma (FSAP-DP) were compared. Substrate turnover was monitored after addition of NNKC9/41 linear or the Ser-Ser control peptide (25 μM each). Results are shown as mean + range of duplicate wells. (B) Pooled pro-thrombin-deficient citrated plasma (1:5 dilution) was recalcified with 10 mM CaCl_2_ and was incubated with NNKC9/41 linear or scrambled control peptide (0-25 μM each) for 60 min at 37°C in the presence or absence of the FSAP-inhibitory MA-FSAP-38C7 (5 μg/ml). Initiation of clotting was achieved by adding tissue factor/phospholipids and prothrombin complex concentrate. Clotting time was measured as diluted prothrombin time (dPT) in s. Data are the mean of 3 independent experiments. The relative dPT in % of the untreated control is presented as mean ± SE (*p <0.05, 2 way ANOVA and Bonferroni post test). Inset shows a magnified *y*-axis.

### Effect of NNKC9/41 on hemostasis

We then addressed the question if the activation of pro-FSAP in plasma with the peptide leads to any biological effects. We have previously shown that FSAP promotes coagulation by inactivating TFPI ^*30*^ so we tested the effect of the peptide on clotting. The use of citrate plasma for this experiment was problematic because the peptide did not function optimally in undiluted citrate plasma and, after calcification, the peptide had a long lag time for pro-FSAP activation whereas clotting is very fast. To overcome these issues we used a clotting assay where the peptide was added to pro-thrombin-deficient plasma in the presence of Ca^2+^ to allow for pro-FSAP activation but without any premature clotting of the plasma. A pro-thrombin containing concentrate was then added to enable tissue factor/phospholipid-triggered clotting. NNKC9/41 had a significant effect on lowering the clotting time and this effect was reversed in the presence of MA-FSAP-38C7 (Fig. 7B). Thus, the NNKC9/41 activated pro-FSAP and MA-38C7 inhibits it, leading to functional consequences for clotting. It should be noted that the antibody also directly inhibits the effect of FSAP on TFPI as shown previously^*30*^.

### Effect of NNKC9/41 on mouse plasma

Mouse FSAP differs in fours amino acids (29-33) from human FSAP in the MA-FSAP-38C7 binding sequence in the NTR (Fig. 3C). This raises the question if the peptide and antibody would function in mouse plasma or not. Fluorescence substrate turnover was low in histone-stimulated wild-type mouse plasma compared to human plasma. There was no substrate turnover in plasma from FSAP^−/−^ mice confirming that substrate turnover was, in fact, due to FSAP and not another protease (Supplementary Fig. S6). The peptide did not activate mouse pro-FSAP and MA-FSAP-38C7 had no effect on histone activation in mouse plasma. Thus, the peptide is likely to be specific for human pro-FSAP only and, in its current form, applications in mouse models *in vivo* are unlikely.

## Conclusions

The activation of the zymogen form of proteases by charged molecules plays a fundamental role in in the mobilization of thrombosis and hemostasis pathways. The contact pathway, which involves the activation of FXII and PK, is mediated by negatively charged molecules ^*31*^. FSAP activation follows a divergent pattern in that it is activated by positively charged macromolecules. In a model of pro-FSAP activation proposed by Yamamichi et al ^*18*^, disruption of a charge-based interaction between NTR and EGF3 domain of pro-FSAP by histones leads to an intermolecular dimerization between two pro-FSAP molecules followed by auto-activation. NNKC9/41, most likely, follows the same mechanism of activation but its effect is not solely charge-driven. At this stage we can say that the peptide, most likely, forms a higher order covalent and/or non-covalent structure that is sequence specific. This peptidic activator of pro-FSAP, as well as its interaction sequence in pro-FSAP, will be a useful tools to establish the molecular details of FSAP zymogen activation. As a corollary, the same information can be used to develop inhibitors of FSAP activation that will help to elucidate the biological role FSAP as well as develop novel therapeutic strategies.

## MATERIALS AND METHODS

### Expanded details of all procedures are in the supplemental section

#### Peptides

Peptides were synthesized by Genescript (Piscataway, NJ, USA) or JPT (Berlin, Germany). For a selection of peptides, the sequence and the number of batches tested in this work is indicated in parentheses in Supplementary Table 1. Where appropriate, the peptides were biotinylated at the N-terminus by the vendor. All peptides were dissolved in DMSO to provide maximal solubility.

### Activation of endogenous pro-FSAP in plasma

Human citrate-plasma (0.38% w/v) or hirudin-plasma (25 ug/ml Lepirudin) was from 5 healthy donors. Plasma was diluted 1:2 - 1:12 and mixed with various test substances that potentially activate or inhibit pro-FSAP. Ac-Ala-Lys-Nle-Arg-AMC (amino-methyl-coumarin) was used as a sensitive and specific substrate for FSAP ^*24*^. Hydrolysis of the fluorogenic substrates was measured using a Synergy HI plate reader with excitation at 320 nm and emission at 460 nm at 37°C for 60 min. The maximal velocity was calculated from the linear part of the progress curve. In limited experiments, the 2^nd^ generation fluorogenic FSAP substrate (Ac-Pro-*D*Tyr-Lys-Arg-AMC)^*32*^ was used. Pro-uPA activation, formation of FSAP-α2-antiplasmin and Western blotting with anti-FSAP antibodies was also used to monitor pro-FSAP activation.

### Statistical analysis

All experiments relating to the activation of purified pro-FSAP or plasma were performed with different batches of synthetic peptides and different donors. Each experiment was performed in duplicates or triplicates and the results are shown as mean ± range or mean ± SEM, respectively. In the plasma clotting assays, results from 3 independent experiments are pooled and are shown as mean ± SEM and the statistical analysis was done using 2-way analysis of variance (ANOVA) followed by Bonferroni test using Graphpad prism.

## Acknowledgements

We would like to thank Jeong Yeon Kim for collecting the mouse plasma, Kristina Byskov for the statistical analysis (both from University of Oslo).

## Author contributions

SS performed the phage screening experiments and the binding studies with peptides. NVN performed all experiments relating to the preparation and characterization of the recombinant proteins and enzyme activity assays. ME performed all the experiments relating to haemostasis. GÅL constructed and provided the phage display libraries as well as the design and analysis of the phage-related experiments. PJD generated and provided the monoclonal antibodies for screening. BEH and BT performed and analysed all the MALDI-TOF and LCMS experiments. MS performed the HPLC. SMK performed all experiments with plasma. SMK designed the study, obtained the funding, analysed the data and wrote the manuscript. All authors edited the final version of the manuscript.

---

**Supplementary Table 1:**
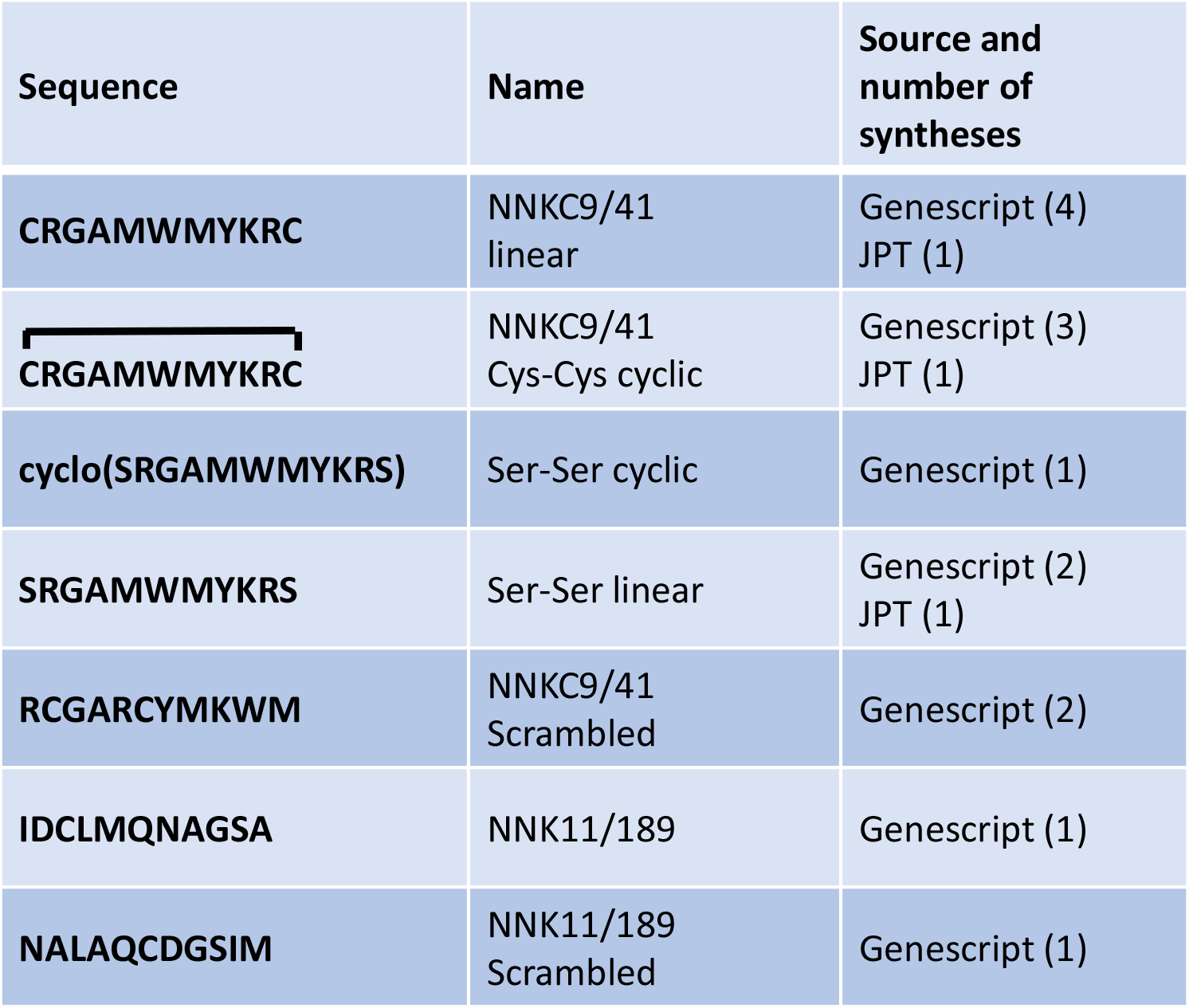
Overview of the peptides synthesized

**Fig S1.**
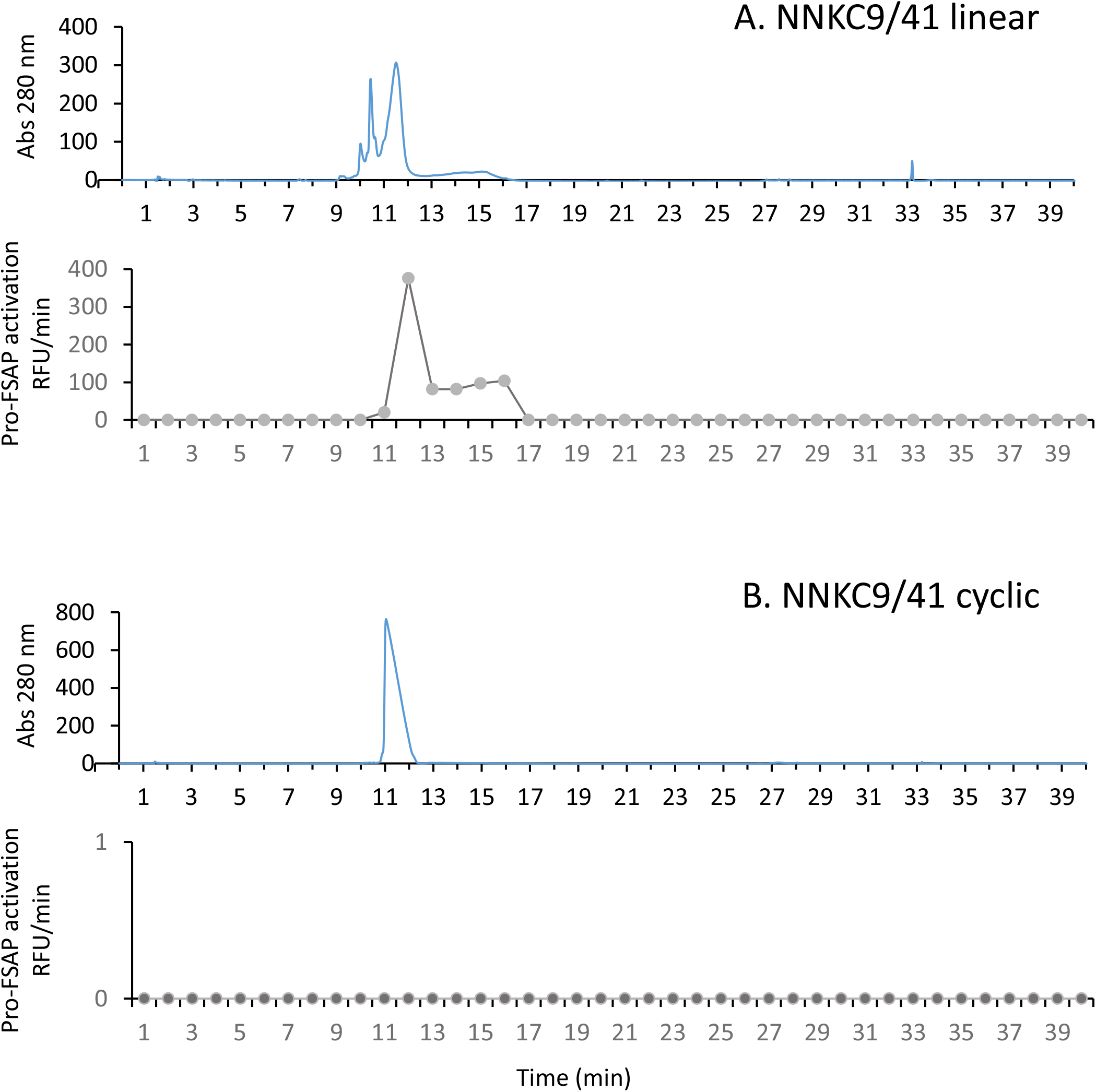
Repurification of synthetic peptides by HPLC and activity profiles of fractions: 100-300 μg each of (A) NNKC9/41 linear and (B) NNKC9/41 cyclic were run on C-18 HPLC. Fractions were collected, lyophilized and reconstituted in 200 μl H_2_0. Hirudin plasma (1:12) was stimulated with 1μl of each fraction and pro-FSAP activation was determined with a fluorescent substrate (Ac-Pro-*D*Tyr-Lys-Arg-AMC, RFU/min).

**Fig S2.**
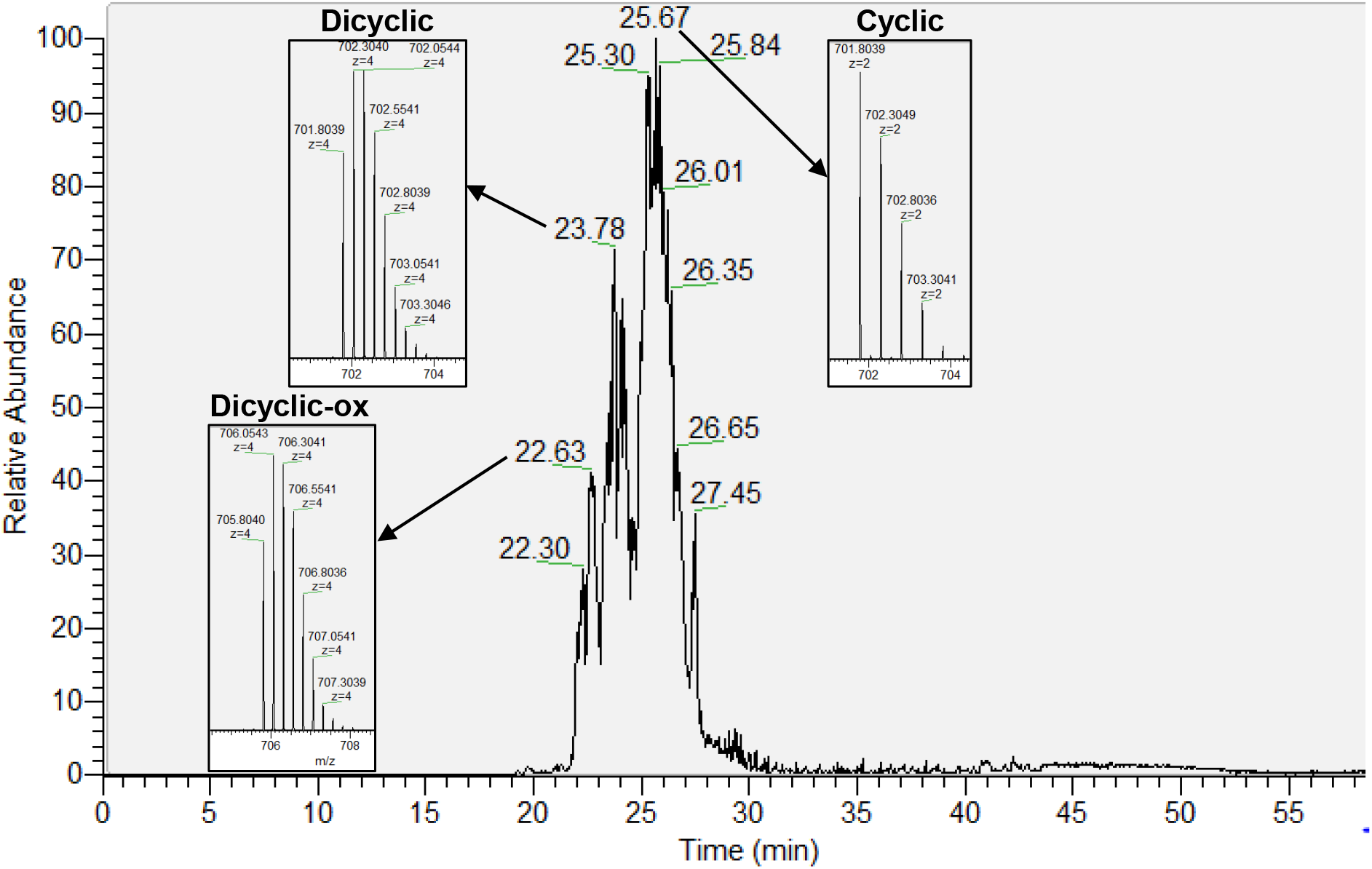
LC-MS analysis of active fractions of HPLC repurified synthetic peptide: The active HPLC fraction according to Fig. S1A (A, fraction 12) was analyzed by LC-MS. The base peak chromatograms are presented and the detected mass-to-charge (m/z) ratios at different peaks were included. The monoisotopic masses (M) corresponded to the molecular masses of NNKC9/41 of the cyclic (m/z=701.80/2; M=1401.6 Da), dicyclic (m/z=701.80/4; M=2803.2 Da), oxidized dicyclic (dicyclic-ox; m/z=705.80/4; M=2819.2 Da), and dioxidized dicyclic (dicyclic-diox; m/z=709.80/4; M=2839.2 Da) form, respectively. Oxidation with oxygen leads to an increase of the molecular mass by 16 Da and is well known to take place at methionines and tryptophanes, respectively.

**Fig S3.**
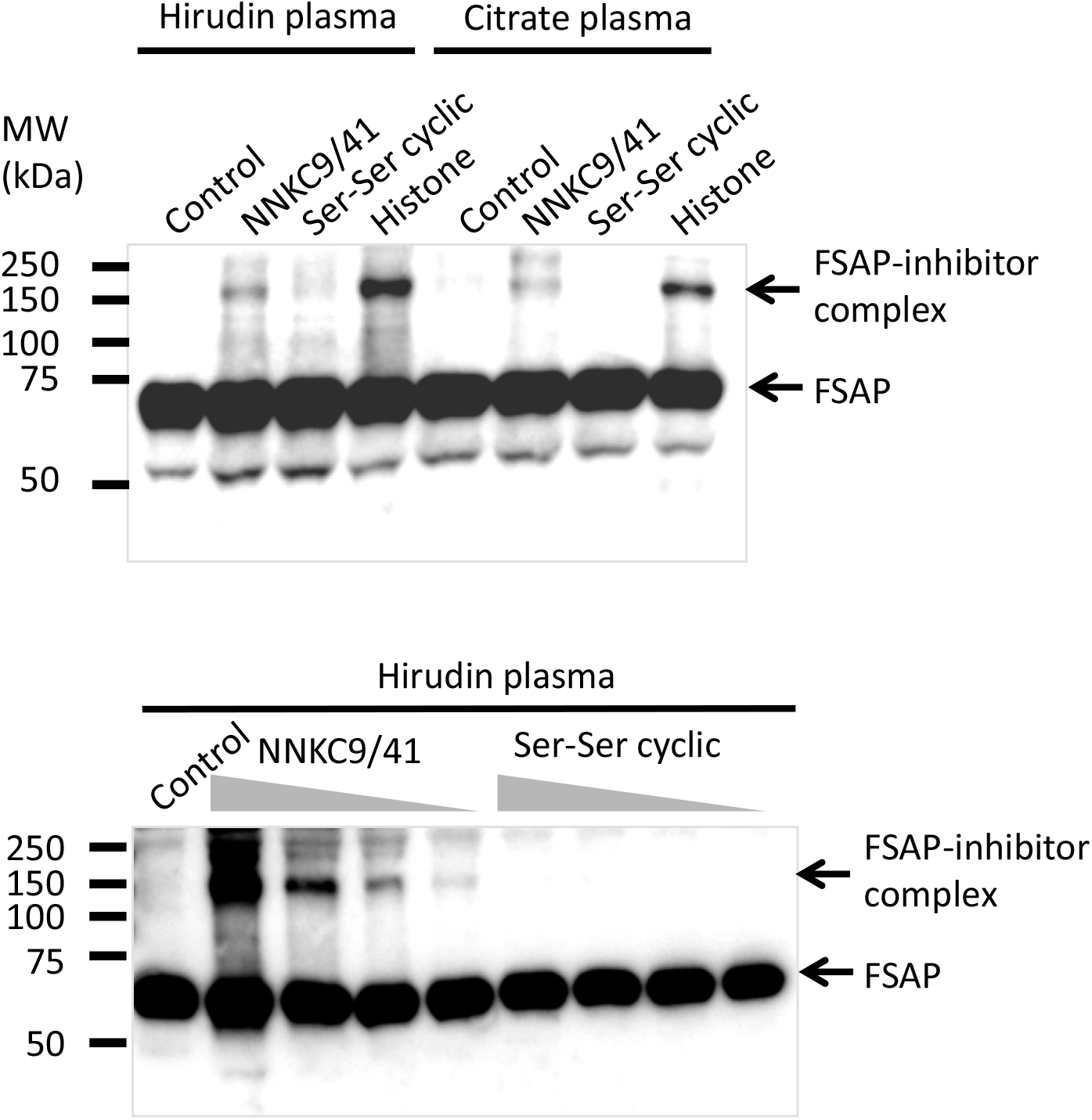
Western blotting of activated plasma: Hirudin or citrate plasma (1:2) dilution was incubated for 1 h at 37 ºC with test substances and the samples were analyzed by Western blotting with an anti-FSAP polyclonal antibody under non-reducing conditions. Top panel: NNKC9/41 linear and Ser-Ser cyclic peptides (25 μM) were compared to histones (50 μg/ml). Lower panel: 100, 50, 25 and 12.5 μM NNKC9/41 linear and Ser-Ser cyclic peptide were compared. Arrows on the right indicate the presence of FSAP and FSAP-inhibitor complexes and MW markers are indicated on the left. Results are representative of 3 independent experiments.

**Fig S4.**
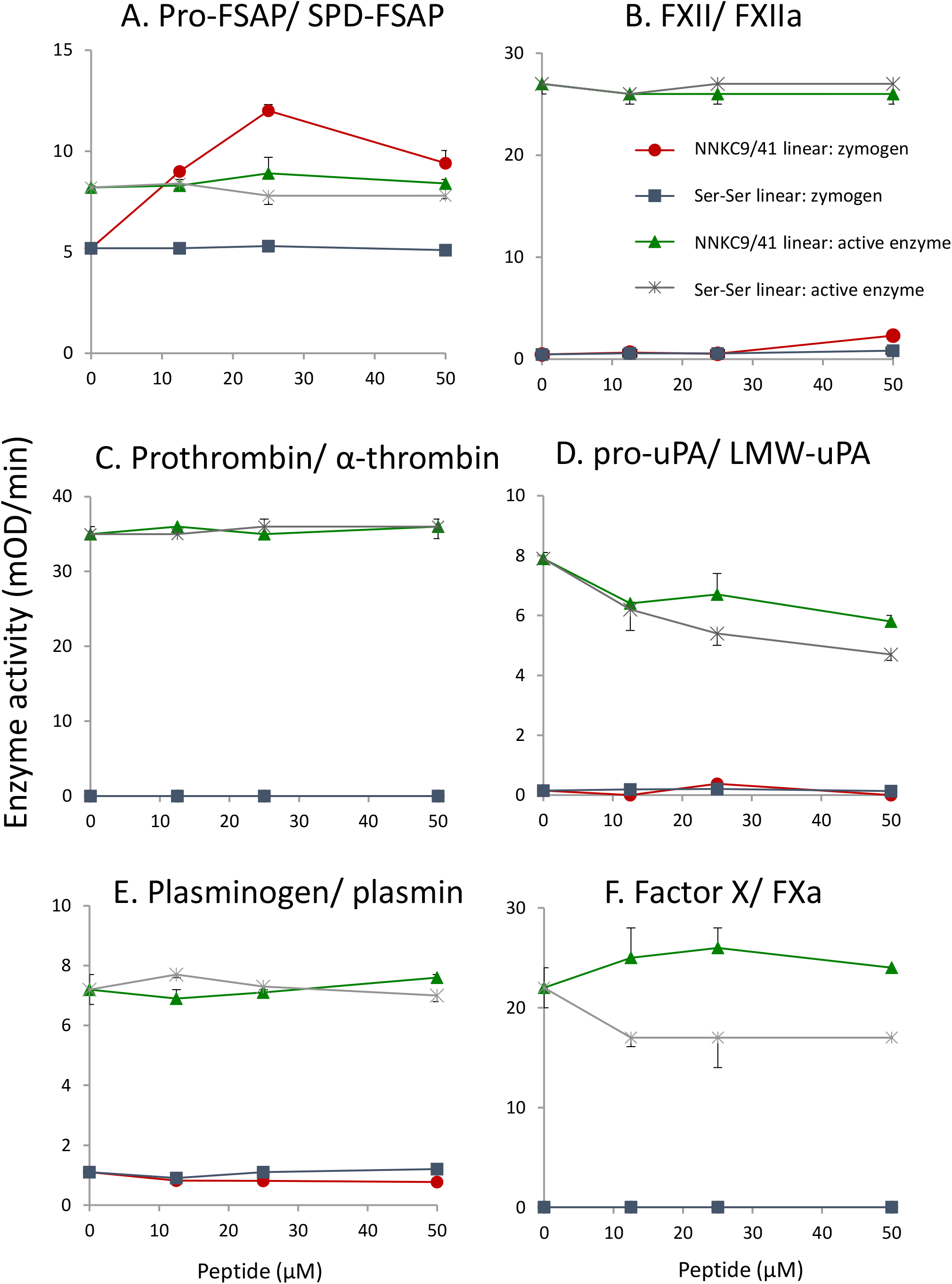
Specificity of NNKC9/41 towards other pro-enzymes: Zymogens and their respective active enzymes were incubated with NNKC9/41 linear or Ser-Ser linear peptide (0-50 μM) and the enzyme activity was monitored using respective chromogenic substrates. Activity of Factor XII (100 nM) and Factor XIIa (20 nM) was measured with the substrate CS31 (1000 μM). Pro-thrombin (100 nM) and thrombin (20 nM) with CS01 (1000 μM). Plasminogen (50 nM) and plasmin (20 nM) with CS41 (1000 μM). Pro-uPA (100 nM) and uPA (20 nM) with PNAPEP 1344 (250 μM). Factor X (50 nM) and Factor Xa (20 nM) with S-2765 (250 μM). pro-FSAP (20 nM) and recombinant serine protease domain (rSPD) (20 nM) with S2288 (250 μM). All pro-enzymes and enzyme were obtained from Enzyme Research Laboratories (South Bend, IN, USA) except plasminogen and FSAP, which were isolated in house; and pro-uPA was from Grünenthal (Stolberg, Germany). All chromogenic substrates were from Hyphen Biomed (Neuville Sur Oise, France). Experiments were carried out in TBS with Tween-20 (0.1 % wt/ vol) BSA (0.3% wt/vol) and CaCl_2_ (2 mM). Absorbance at 405 nm was measured over time and the enzyme activity is depicted was measured in duplicates (mean + or - range) and similar results were obtained in 3 independent experiments.

**Fig S5.**
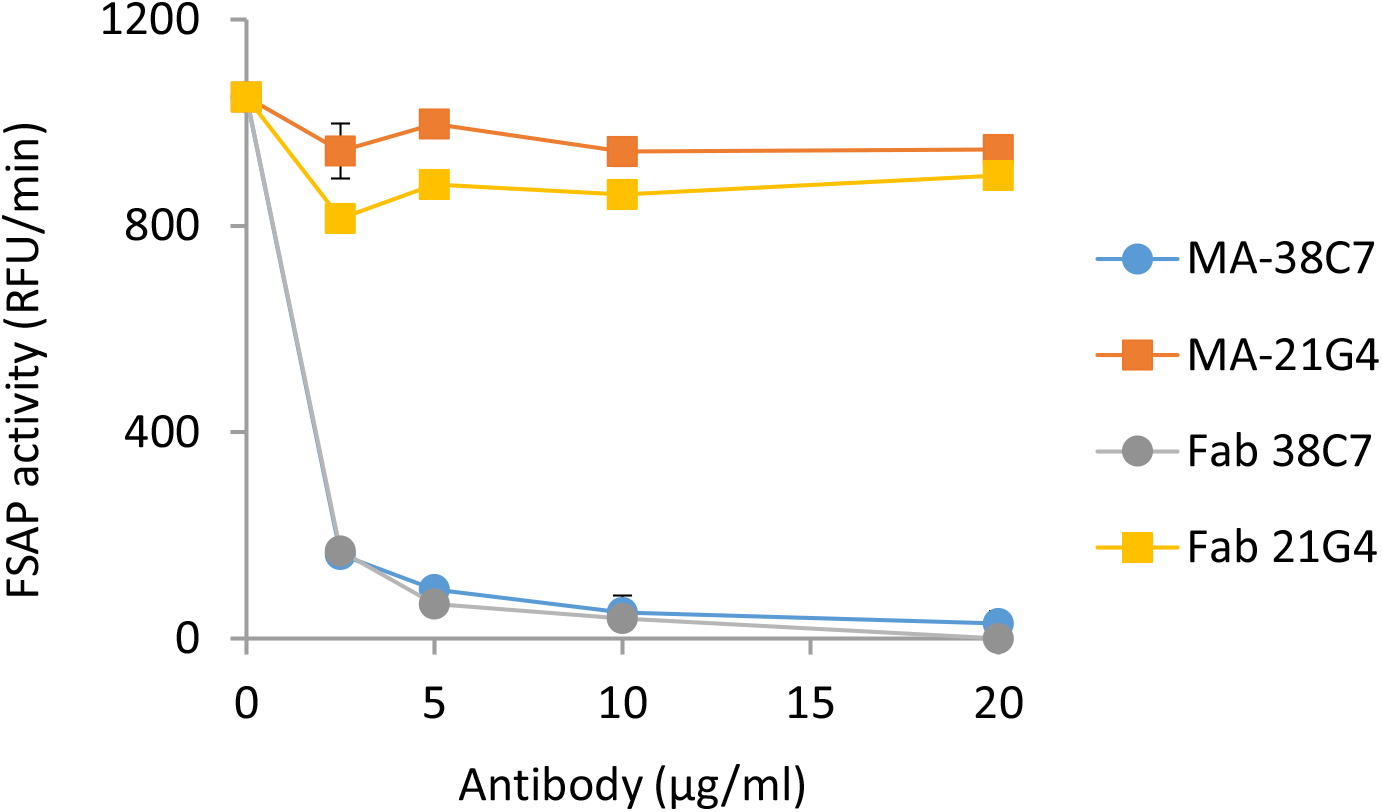
Effect of anti-FSAP antibodies on histone-mediated pro-FSAP activation in plasma: Hirudin plasma was stimulated with histone (25 μg/ml) in the presence of increasing concentration of MA-FSAP-38C7 and a control antibody (MA-FSAP-21G4) or their Fab fragments as indicated. Turnover of the FSAP fluorescent substrate (Ac-Pro-*D*Tyr-Lys-Arg-AMC) was measured in duplicate (mean +/- range). The antibody and Fab fragments were compared on a weight basis and not a molar basis. Similar results were obtained in 2 independent experiments.

**Fig S6.**
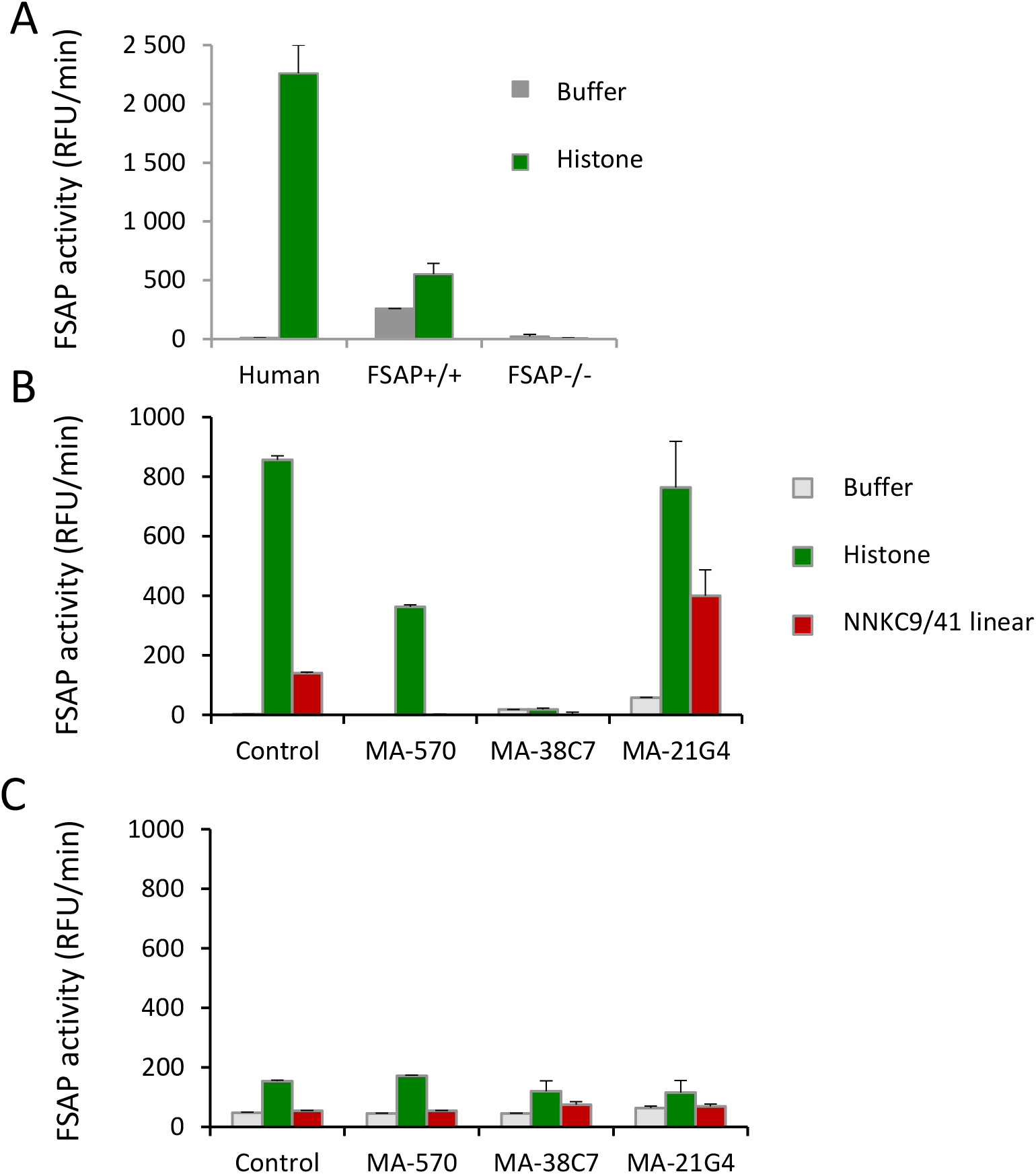
Effect of NNKC9/41 and MA-FSAP-38C7 on human and mouse plasma: (A) Human citrate-plasma and mouse citrate plasma from FSAP^+/+^ or FSAP^−/−^ mice was stimulated with histone (25 μg/ml). (B) Human citrate-plasma and (C) mouse citrate plasma was stimulated with histone (25 μg/ml) or NNKC9/41 linear (40 μM) in the presence of MA-FSAP-570, MA-FSAP-38C7 or MA-FSAP-21G4 (20 μg/ml each). Plasma was diluted 1:10 and turnover of the FSAP fluorescent substrate (Ac-Pro-*D*Tyr-Lys-Arg-AMC) was measured in duplicate (mean + range). See also sequence comparison between mouse and human in the NTR of FSAP in Fig. 3C which shows a difference in the presumed peptide binding sequence. Similar results were obtained in 2 independent experiments.

## SUPPLEMENTARY MATERIALS AND METHODS

### Helper phages, bacterial strains and construction of peptide phage libraries

The M13K07 was purchased from GE Healthcare (Uppsala, Sweden), whereas Delta Phage was prepared as described ^1^. The *E. coli* strains SS320 and XL1-Blue were purchased from Lucigen Corp. (Middleton, WI, USA) and Life Technologies (Carlsbad, CA, USA), respectively. A library template phagemid was prepared by inserting a short oligo containing two consecutive stop codons (ochre/opal) into the *Nco*I/*BamH*I expression cassette of the pGALD9ΔL phagemid ^2^ using standard methods. Using this template, a linear random 11-mer (NNK11) and a Cys-constrained random 9-mer (NNKC9) peptide library fused to pIX was constructed based on Kunkel mutagenesis essentially as described ^3^. The final *E. coli* SS320 transformation frequency-based library sizes were about1 × 10^10^ unique clones and the libraries were prepared high valency peptide display by rescue with DeltaPhage.

### Phage selection with human FSAP

Tubes and streptavidin-coated Dynabeads (Thermofischer Scientific, Oslo, Norway) were blocked with PBS with 4% (w/v) skim milk powder. Approximately 1 × 10^13^ virions of library and 1-10 μg (R1: 10 μg, R2 – 3: 1 μg) of biotinylated pro-FSAP were incubated with the appropriate amount of Dynabeads for 60 min at RT. The beads/protein/phage complexes were separated from the supernatant using a magnetic rack and washed with PBS-T20 (0.1% (v/v) Tween-20 in PBS). The complexes were then incubated with an excess amount of unbiotinylated pro-FSAP for 60 min at RT to remove phages bound with low affinity and retain only high affinity binding clones. The beads/protein/phage complexes were then separated from the supernatant containing the soluble FSAP using a magnet rack and washed with PBS-T20 (0.1% (v/v) Tween-20 in PBS) and with PBS pH7.4. This was applied to both libraries and in all rounds. The remaining phages were then eluted from the beads with 500 μL of 100 mM triethylamine (pH 11) and neutralized in Tris-HCl pH 7.4.

### Bacteria preparation, library amplification and phage particle preparation

*E. coli* XL1-Blue bacteria were grown in 2xYT-TAG (30 μg/ml tetracycline, 100 μg/ml ampicillin and 0.1M glucose) medium at 37°C overnight and use to make a fresh rescue culture. Phages eluted from R1 were added to log-phase cells in 2xYT-TAG, followed by incubation at 37°C. After centrifugation, the pellet was re-suspended, and the solution was plated onto 2x YT-TAG Q-trays and incubated at 30°C overnight. The next day, cells were scraped from the agar plates with a total volume of 20 ml 2xYT. The scraped material was used to re-inoculate 50ml 2x YT-TAG medium at 37 °C until it reached an OD600 of 0.2 before super-infection (multiplicity of infection 20) with M13K07 helper phage. Cells were pelleted by centrifugation and the pellet was gently re-suspended in 50 ml pre-warmed 2xYT AK (100 μg/ml ampicillin and 50 μg/ml kanamycin) and incubated at to produce phage particles displaying peptides at low valence. The culture was centrifuged at 10,000 rpm for 10 min and the supernatant was sterile filtrated using 0.2μm filter. The phage particles were purified and concentrated by PEG/NaCl precipitation as described ^4^. Virion concentration was determined by the formula: virions/ml=[(A_269nm_-A_320nm_) × 6.083×10^16^]/ genome size ^5^.

### Binding studies with biotinylated peptides

Wells were coated with anti-FSAP polyclonal antibody (5 μg/ml). Plates were washed with TBS (25 mM Tris-HCl (pH 7.5) and 150 mM NaCl) containing 0.2 % (w/v) Tween 20 (TBS-T) and were blocked with TBS containing 3% (w/v) BSA (Sigma). Recombinant FSAP proteins were captured and biotinylated peptides were added to allow binding. Subsequently, the plates were washed and bound peptides detected with peroxidase-coupled streptavidin. The non-specific binding to BSA was subtracted from binding to test protein to calculate specific binding.

### Recombinant protein expression

FSAP cDNA was derived from human liver RNA and subsequently cloned into the pASK-IBA33plus vector (IBA-lifesciences, Goettingen, Germany). This vector contains a C-terminal 6xHIS tag. The construct of the recombinant N-terminal FSAP comprising amino acids 24-313 was used as template to generate mutants with the following amino acids deletions; ΔNTR (24-72), ΔEGF1 (73-109), ΔEGF2 (111-148), ΔEGF3 (150-188), ΔKringle (193-276) respectively. The serine protease domain construct comprises amino acids 292-560 and its expression has been described before ^6^.

Proteins were expressed in BL21-Gold (DE3) in inclusion bodies. Washed inclusion bodies were resuspended in 100 mM Tris, 150 mM NaCl, 5 mM 2-mercaptoethanol, 8 M urea pH 8 for 1h followed by a centrifugation at 14000 rpm for 10 min. After purification on a Ni-NTA-Agarose column the protein was diluted to a concentration of 0.1mg/ml and dialyzed in 100 mM Tris, 150 mM NaCl, 5 mM 2-ME, 8 M urea pH 8 for 1h at room temperature. This was followed by dialysis in 20 mM Tris, 10% (v/v) glycerol 4 M urea, pH 8 at 4 °C overnight and finally in 20 mM Tris pH 8, 10% (v/v) glycerol overnight at 4 °C. The protein was cleared for any precipitation and concentration was determined by Bio-Rad reagents using BSA as a standard. Protein quality was checked by Coomassie Blue staining of SDS-PAGE gels.

### Pro-FSAP enzyme activity assay

Pro-FSAP was isolated from human plasma as described before^7^. The content of the zymogen form varied from 50-90% due to activation during the purification process. Histones, isolated from calf thymus, were obtained from Sigma Aldrich (Oslo, Norway). FSAP activity assays were performed as described previously ^8^. In brief, microtiter wells were blocked with TBS (25 mM Tris-HCl, pH 7.5, and 150 mM NaCl) containing 3% (w/v) BSA for 1 h and washed with TBS-T. The standard assay system consisted of TBS-T with 0.3 % (w/v) BSA and CaCl_2_ (2 mM) 1 μg/ml (15 nM) plasma purified pro-FSAP and 250 μM chromogenic substrate S-2288 (D-Ile-L-Pro-L-Arg-*p*-nitroaniline dihydrochloride) (Haemochrome, Essen, Germany) and was followed at 37°C at 405 nm in a microplate reader Synergy HI plate reader (BioTek Instruments, Winooski, USA). In some experiments, a fluorescent substrate of FSAP^9^ was used as described below.

### Effect of FSAP-activating peptide NNKC9/41 on the extrinsic pathway of coagulation

Pro-thrombin-deficient citrated pool plasma (Haemochrom Diagnostica, diluted 1:5 in HBS, pH 7.4) was recalcified with 10 mM CaCl_2_ and was incubated with peptides for 60 min at 37 °C in the presence or absence of the FSAP-inhibitory MA-FSAP-38C7 (5 μg/ml). Initiation of clotting was achieved by adding 100 μl tissue factor/phospholipids, Thromborel (Siemens Healthcare, Marburg, Germany), 1:3000 prediluted in HBS and, prior to use, adjusted to 0.3 IU/ml FII by addition of a prothrombin complex concentrate (Ph. Eur. quality). Final concentrations in the clotting assay were 1:8 diluted plasma, 1:7500 Thromborel, 0.125 IU/ml prothrombin. Clot formation was measured optically at 405 nm in a platereader (Tecan, Crailsheim, Germany). The time from starting the reaction to 50 % maximum clot turbidity was measured as diluted prothrombin time (dPT) in s.

### HPLC

The HPLC separation of the peptides was carried out on a ZORBAX 300SB-C18 column (4.6mm i.d x 150mm, 5-micron, Agilent) and UV detection at 220nm and 280nm. 100-300μg peptide was injected. The flow rate used was 1mL/min, and the solvent gradient was 5% to 65% B in 25 minutes and then to 95% B in 6 minutes. Solvent A was aqueous 0.065 % trifluoroacetic acid, whereas solvent B was 100 % acetonitrile in 0.05 % trifluoroacetic acid. Time-based fractions (1 min) were collected and dried in a speed vac.

### LC-MS

The HPLC purified fractions were dried using a Speed Vac concentrator (Concentrator Plus, Eppendorf, Hamburg, Germany), dissolved in 10 μl 0.1% formic acid/2% acetonitrile and 5 μl analyzed using an Ultimate 3000 RSLCnano-UHPLC system connected to a LTQ Oribtrap XL mass spectrometer (Thermo Fisher Scientific, Bremen, Germany) equipped with a nanoelectrospray ion source. For liquid chromatography separation, an Acclaim PepMap 100 column (C18, 2 μm beads, 100 Å, 75 μm inner diameter, 50 cm length) (Dionex, Sunnyvale CA, USA) was used. A flow rate of 300 nL/min was employed with a solvent gradient of 4-35% B in 60 min. Solvent A was 0.1% formic acid and solvent B was 0.1% formic acid/90% acetonitrile. The mass spectrometer was operated in the data-dependent mode to automatically switch between MS1 and MS2 acquisition. Survey full scan MS spectra (from m/z 300 to 2,000) were acquired with the resolution R = 60,000 at m/z 400 (LTQ-Orbitrap XL), after accumulation to a target of 1e6. The maximum allowed ion accumulation times were 60 ms. The method used allowed sequential isolation of up to six most intense ions, depending on signal intensity (intensity threshold 1.7e4), for fragmentation using collision induced dissociation (CID) at a target value of 10,000 charges in the linear ion trap of the LTQ-Orbitrap XL. Target ions already selected for MS2 were dynamically excluded for 60 sec. For accurate mass measurements, the lock mass option was enabled in MS mode. Data were acquired and analyzed using Xcalibur v2.5.5.

